# ReScale4DL: Balancing Pixel and Contextual Information for Enhanced Bioimage Segmentation

**DOI:** 10.1101/2025.04.09.647871

**Authors:** Mariana G. Ferreira, Bruno M. Saraiva, António D. Brito, Mariana G. Pinho, Ricardo Henriques, Estibaliz Gómez-de-Mariscal

**Author notes:** Correspondence |.

## Abstract

Deep learning has established itself as the state-of-the-art approach for segmentation in bioimage analysis. However, these powerful algorithms present an intriguing paradox regarding image resolution: contrary to intuition, lower-resolution images can yield superior performance for specific image analysis carried out by deep learning. This phenomenon is particularly significant in microscopy imaging, where high-resolution acquisitions come with substantial costs in throughput, data storage requirements, and potential photodamage to specimens. Through systematic experimentation, we evaluate how varying image resolution impacts deep learning performance in cellular image segmentation tasks. We trained popular architectures on datasets downsampled to 6-50% of their original resolution, mimicking acquisitions at lower image magnification, and compared their performance against models trained on native-resolution images. Our results show that segmentation accuracy either improves (by up to 25% of mean Intersection over Union (IoU)) or experiences only minimal degradation (< 5% of mean IoU) when using images downsampled by up to a factor of 4 (25% of the original resolution). This downsampling proportionally increases information throughput while reducing data storage requirements and inference time. With these findings, we contribute systematic guidelines to deep learning practitioners in creating efficient experimental pipelines for image-driven discoveries. This approach improves the sustainability and cost-effectiveness of bioimaging studies by reducing data and computing needs while optimising microscopy techniques.

## Introduction

Modern microscopy technologies have driven an unprecedented expansion in data volume. High-resolution techniques, particularly super-resolution microscopy (1), generate substantially larger datasets than traditional methods when imaging equivalent sample areas. A single 3D multichannel cell image from a spinning disk confocal requires approximately 1GB of storage (considering a size of 4MB for each slice of a three-channel and 100 *z*-slices volume). In contrast, the same field acquired with a super-resolution laser scanning microscope would be approximately 60 times larger (estimated from images acquired in-house). To address the challenges in acquisition, storage, and processing posed by these volumes of data, researchers increasingly rely on deep learning approaches that automate analytical tasks while delivering reliable and reproducible results. For this, the bioimage analysis community has developed numerous user-friendly solutions (2, 3), including standalone software and platform-integrated plugins (4–11), enabling researchers to leverage these powerful methodologies.

Yet, bioimage analysis faces unique efficiency challenges compared to other deep learning domains. The most significant bottlenecks include: limited computational resources and storage capacity, particularly problematic for time-lapse volumetric data; scarcity of large annotated datasets essential to train robust models and transfer learning (12, 13); and insufficient understanding of the relationship between image quality metrics and computational performance requirements (14, 15).

In computer vision, image quality often equates to an algorithm’s efficiency in extracting relevant information under hardware constraints. Counterintuitively, a low-resolution image containing sufficient detail to perform the computational task accurately may be optimal, as it maximises the information-to-resource ratio. For instance, cell nuclei appear more homogeneous at lower resolutions, enabling neural networks to generalise more effectively. Conversely, at high resolution, nuclei images can display heterogeneous structures, introducing noise during segmentation and potentially increasing false negatives. This computational perspective contrasts with traditional microscopy practices, where image quality has been equated with higher resolution and magnification, assuming finer details provide more informative content (1). However, while visually appealing, the wealth of information in high-resolution images can be irrelevant or even detrimental to computational analysis, as our results here demonstrate.

We present this phenomenon as the **image resolution paradox**: While high-resolution microscopy captures more detail, this additional data may contribute minimally—or even negatively—to a certain analysis task while requiring substantially greater resources. This paradox is particularly evident in deep learning-based pipelines (Fig. 1). The limitation stems from convolutional neural network’s (CNN) receptive fields—the spatial region influencing a single output pixel. When image resolution exceeds the network’s architectural optimum, pixels lack sufficient contextual information, compromising the model’s performance. Although practitioners often address this by downsampling during pre-processing, incorporating resolution considerations directly into the experimental design would enable acquisition at optimal resolutions from the outset, simultaneously increasing throughput and reducing computational demands.

**Fig. 1.**
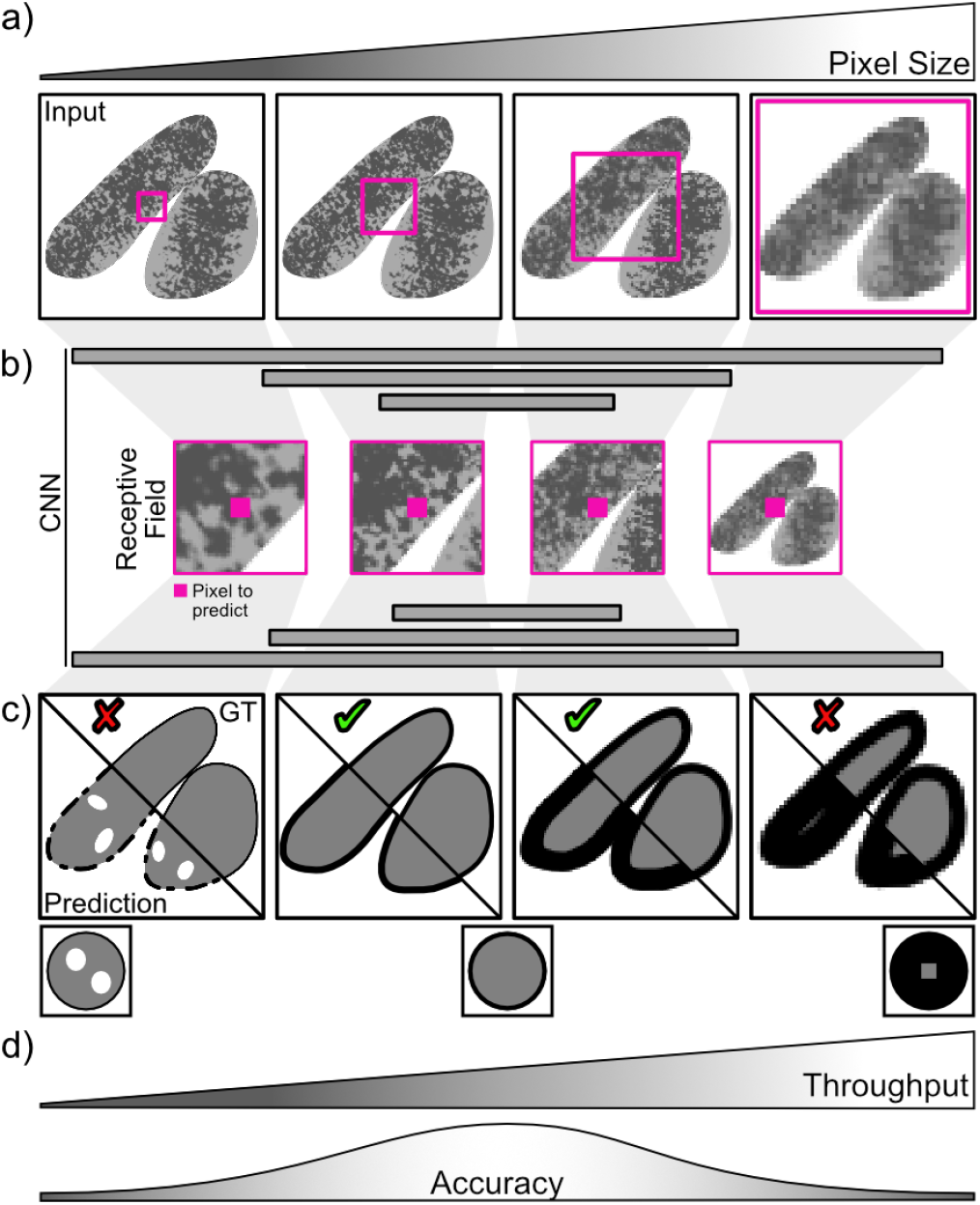
The Image Resolution Paradox. The counterintuitive relationship between image resolution and deep learning performance emerges from the mismatch between pixel size and the network’s receptive field. a) Decreasing image resolution (increasing pixel size) causes pixelation and limits observable details, yet may improve contextual information. b) The area observable within a convolutional neural network’s (CNN) receptive field (magenta squares) determines the context available for predicting the value of a single pixel. c) Suboptimal pixel sizes lead to impaired predictions—either fragmented segmentations caused by non-continuous edges and inner pixels misclassified as background, referred to as false negatives (too high resolution) or over-generalisation at edges (too low resolution). d) Optimal image resolution for CNN-based processing depends on object size rather than maximal achievable microscopy resolution, and proper calibration can simultaneously enhance segmentation accuracy and experimental throughput.

While this paradox is familiar to computer vision experts, few tools explicitly account for it. Cellpose (16) represents an exception, implementing automated re-sampling to optimise pixel-to-cell-diameter ratios. However, most users remain unaware of this critical consideration, leading to sub-optimal application of deep learning methods—particularly with architectures like vanilla U-Net—and unnecessary prioritisation of high-resolution acquisition despite its potential drawbacks.

In this study, we quantitatively demonstrate this paradox and its impact on segmentation performance in bioimage analysis. For clarity, we define image resolution as the information content per pixel, separate from optical resolution considerations. We systematically evaluate resolution effects on model accuracy using a canonical 2D U-Net (17) for semantic segmentation and StarDist (9) for instance segmentation. Our experiments employ diverse specimens, including high-magnification *Caenorhabditis elegans* (*C. elegans*) (18) and brightfield *Escherichia coli* (*E. coli*) (19) images. Additionally, we analyse *Staphylococcus aureus* (*S. aureus*) imaged with Structural Illumination Microscopy (SIM), demonstrating that critical phenotypic changes remain detectable even when reducing resolution by 16*×*, enabling proportional throughput improvements. Beyond these examples, we provide a generalisable framework, ReScale4DL (https://github.com/HenriquesLab/ReScale4DL), that researchers can implement to determine optimal pixel sizes for their specific contexts, maximising both computational efficiency and information extraction (Fig. 2).

**Fig. 2.**
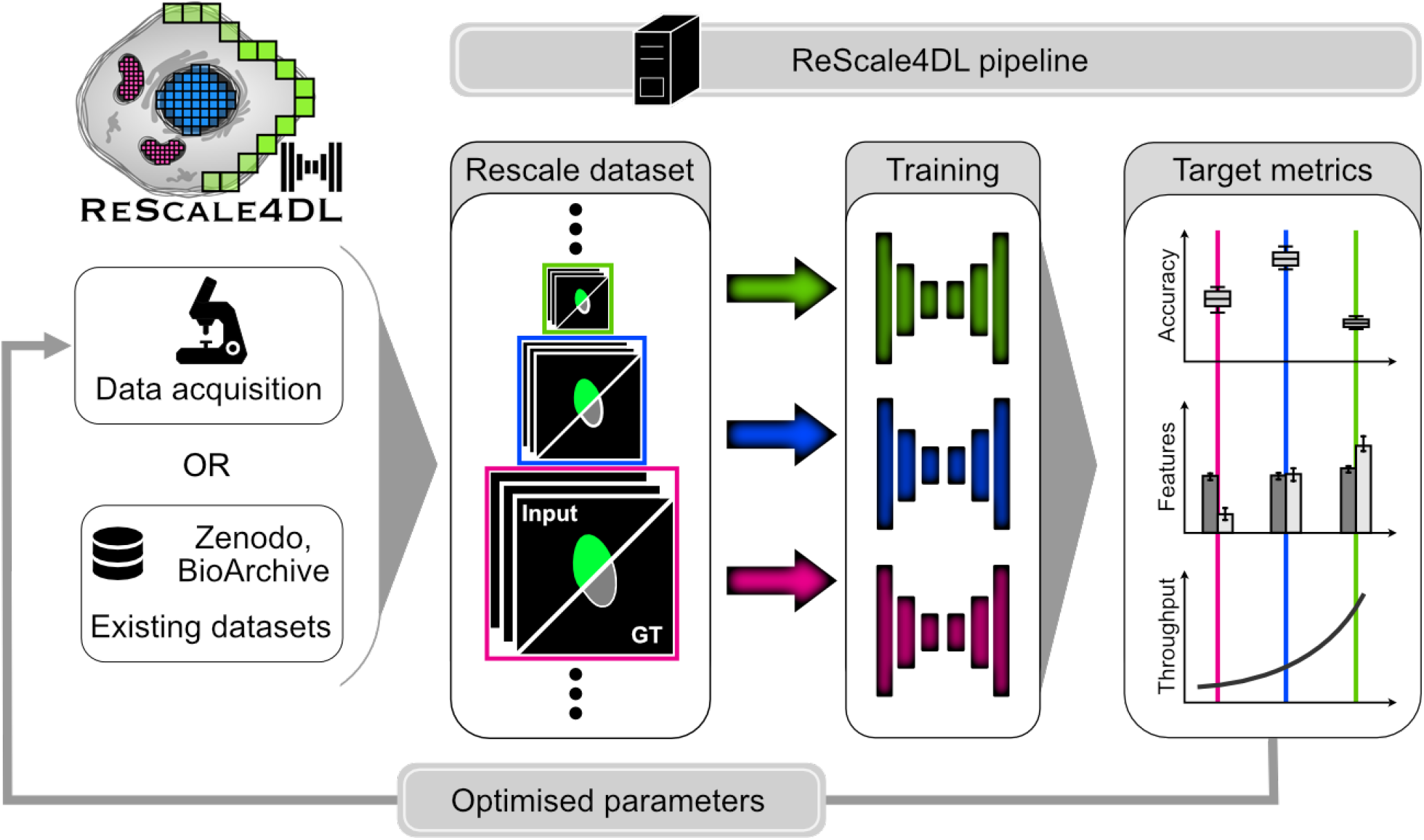
ReScale4DL pipeline. Researchers can optimise the pixel size for a chosen deep learning architecture using their newly acquired and annotated images or on existing datasets, which can be used as a starting point for the optimisation. A series of image rescaling factors is applied to the data, a deep learning model is trained for each rescaled dataset, and different accuracy metrics and features are computed. Once an ideal target metric is chosen, such as the accuracy of the image processing performance, the recovery of an image feature, or a good compromise between the throughput and accuracy, researchers can identify an adequate pixel size and launch their imaging experiments with optimised image resolution. The results presented in this manuscript were obtained by following the ReScale4DL pipeline.

## Results

### High resolution and segmentation accuracy: a counter-intuitive relationship

We trained a 2D U-Net architecture to semantically segment different cellular regions - inner area, edges, and background - using two distinct datasets: *C. elegans* captured with brightfield microscopy and *E. coli* imaged via phase contrast microscopy. For each dataset, we systematically rescaled the images to multiple resolutions and trained separate U-Net models to evaluate performance across these conditions (Fig. 2). Since segmentation accuracy depends on the relationship between image resolution and object size, we quantified the performance relative to the percentage of an object’s diameter covered by a single pixel. Our experiments revealed a counterintuitive relationship between resolution and segmentation quality (Fig. 3). Contrary to conventional expectations, high-resolution images often produced fragmented segmentation masks with unexpected discontinuities. In the *C. elegans* dataset, the inner masks produced at the original resolution contained spurious holes where pixels were misclassified as background (Fig. 3a)). This occurred because the model lacked sufficient contextual information to confidently identify inner and outer objects’ areas and one-pixel width edges were under-represented for the given receptive field and resolution. Namely, the Intersection over Union (IoU) of semantic segmentations decreased by approximately 40% compared to binary segmentation, which remained relatively high (*>* 0.9) despite these deficiencies (Fig 3b)). Notably, downsampling the images improved accuracy metrics for both binary and semantic segmentation approaches, providing clear evidence that maximum resolution does not necessarily yield optimal segmentation results. The *E. coli* dataset analysis corroborated these findings (Fig. 3c-e)).

**Fig. 3.**
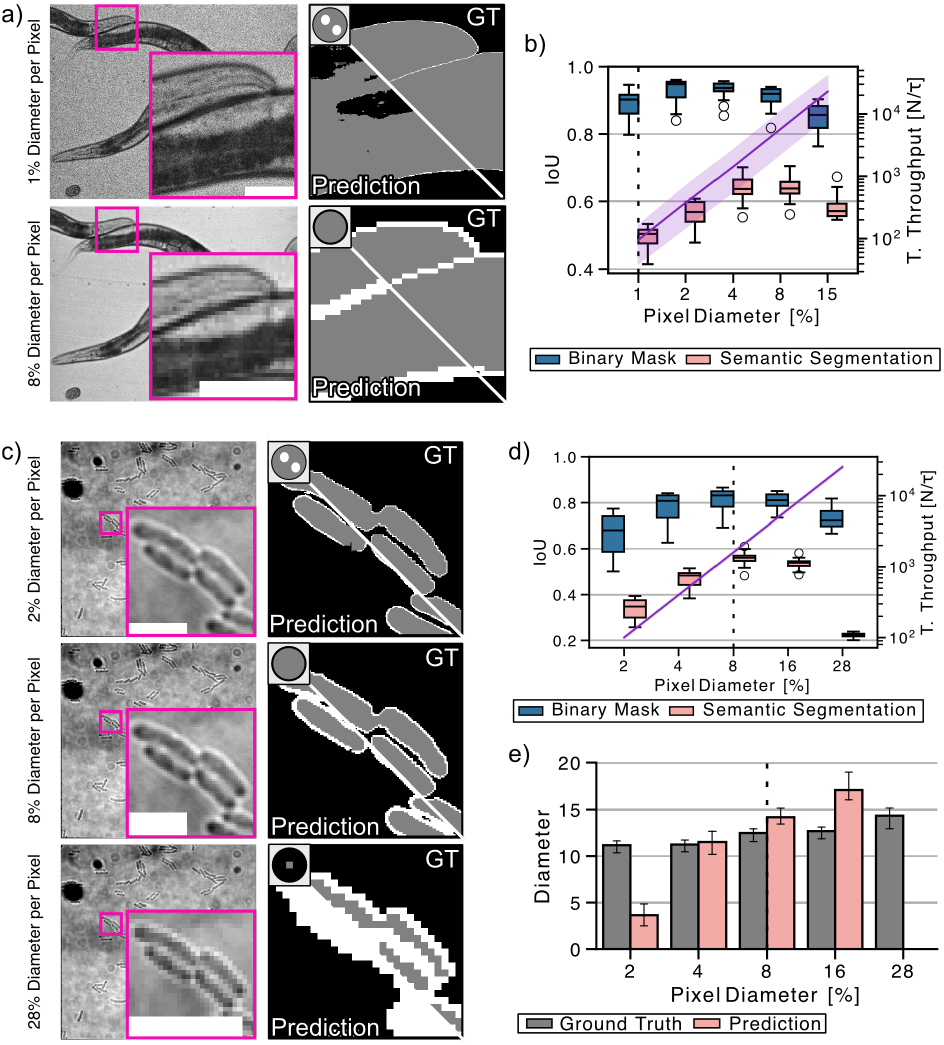
Impact of image resolution on deep learning segmentation accuracy. A 2D U-Net architecture was employed for semantic segmentation of cellular regions (inner area and boundary) in microscopy images. Images were systematically resized to simulate varying acquisition resolutions, with a separate U-Net model trained for each resolution factor. a) Segmentation results for *Caenorhabditis elegans* (*C. elegans*) using original images (pixel covering *∼*1% of worm diameter) and images downsampled by factor 2^3^ (pixel covering *∼*8% of worm diameter). Scale bars: 100 and 25 pixels, respectively. b) (Boxplots) Intersection over Union (IoU) distributions across downsampling factors ranging from *×*1 to *×*2^4^, showing how segmentation accuracy varies with pixel size relative to object diameter. IoU was calculated both for semantic segmentation (average IoU of edge and inner regions) and binary masks (merged labels). (Purple line in logarithmic scale) Theoretical throughput was calculated as the number of objects observable by a unit of time (*τ*) given a certain resolution. *τ* is specific to each microscopy acquisition modality and preserved across the different resolutions. c-e) Equivalent analysis for *Escherichia coli* (*E. coli*) segmentation in phase contrast microscopy. c) Examples showing fragmented results (2% diameter coverage), optimal segmentation (8% coverage), and edge over-segmentation. Scale bars: 100, 25, and 10 pixels, respectively. d) IoU distribution (boxplots) and theoretical throughput (purple line in logarithmic scale) across varying bacteria diameter coverage ratios. e) Comparison between diameter distributions estimated from U-Net segmentation versus ground truth, with closest correspondence at 4% diameter coverage. Vertical dashed lines in panels b), d), and e) indicate the original dataset resolution.

To ensure consistent comparison, we maintained identical U-Net architecture and hyperparameters across all experiments in Fig. 3, enabling direct assessment of resolution’s impact on model performance. Both datasets’ accuracy curves exhibited similar quadratic relationships between image resolution, object size, and IoU scores. For *C. elegans*, binary and semantic segmentation achieved peak IoU when pixels covered 2-4% and 4-8% of worm diameter respectively, demonstrating that optimal resolution differed from the original microscopy acquisition (1% coverage). Similarly, the *E. coli* dataset exhibited the highest IoU at comparable proportional resolutions: 4-16% diameter coverage per pixel for binary segmentation and 8-16% for semantic segmentation. This consistency across different biological specimens reveals a fundamental principle: our U-Net’s 50 *×* 50 pixel receptive field imposes practical constraints. Objects with diameters exceeding 25 pixels (corresponding to < 4% diameter coverage per pixel) become problematic as their size exceeds the receptive field, while objects smaller than 6 pixels (> 16% diameter coverage) risk information dilution and loss through the network’s operations. As exemplified, for any given CNN architecture, there exists an optimal resolution that maximises segmentation accuracy—and importantly, this resolution is typically significantly lower than what human observers prefer, allowing imaging pipelines with substantially improved throughput.

### Optimising image resolution preserves phenotypic discrimination while maximising experimental throughput

Building upon our findings, we demonstrate how optimising image resolution preserves segmentation performance while enhancing experimental throughput. We tested this approach using *Staphylococcus aureus* (*S. aureus*) bacteria treated with antibiotic PC190723 and imaged via SIM. PC190723, a FtsZ inhibitor, prevents cell division and produces enlarged bacteria (20). To quantitatively assess this morphological effect, we employed StarDist—a deep learning approach optimised for instance segmentation of star-convex objects like round bacteria—to segment individual cells and measure diameter distributions (Fig. 4a)). As in our previous experiments, IoU values across different resolution factors revealed an optimal range where segmentation accuracy peaks. Notably, even at substantially reduced resolutions, StarDist maintained robust performance: with up to 11% of bacterial diameter covered by a single pixel, IoU values exceeded 0.8, and even with 23% coverage, IoU remained above 0.75 (Fig. 4b)). Analysis of wild-type (WT) diameter distributions confirmed that measurements derived from ground truth and StarDist predictions remained consistent across resolutions ranging from 0.75% to 6% of diameter per pixel. Only at coverage ratios exceeding 11% (downsampling factor > 2^3^) did diameter estimates begin to diverge, likely because objects were reduced to just a few pixels in width (Fig. 4c)). Most importantly, when training a StarDist model with WT and antibiotic-treated mixed bacteria populations, the diameter differences remained distinguishable across all sampling ratios, demonstrating that phenotypic discrimination is preserved even at significantly lower resolutions than typically employed (Fig. 4d)). Thus, by strategically reducing acquisition resolution, researchers can substantially increase imaging throughput and decrease acquisition times (Figs. 3b), 3d) and 4b)), enabling more comprehensive dataset collection for statistically robust experimental analyses.

**Fig. 4.**
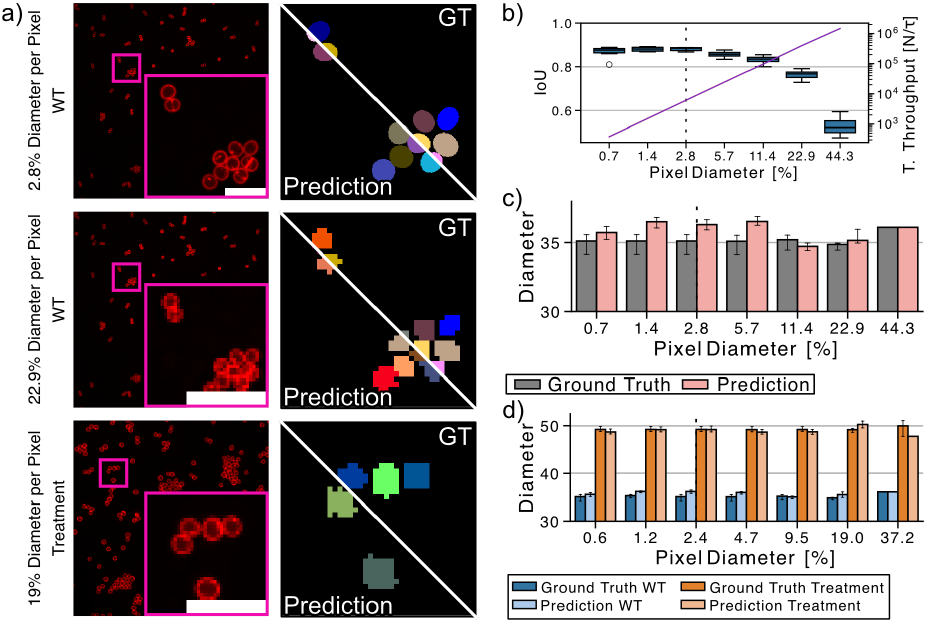
Deep learning-driven resolution adjustment enables phenotype distinction while increasing imaging throughput. *Staphylococcus aureus* (*S. aureus*) bacteria were treated with FtsZ inhibitor PC190723, which prevents cell division and results in larger bacteria phenotypes (20). *S. aureus* (wild type (WT) and treated) were then imaged using Structured Illumination Microscopy (SIM) and segmented with StarDist. a) Representative images of *S. aureus* with ground truth (GT) segmentation and StarDist results at the original resolution (WT) and after 2^3^-factor downsampling (both WT and treated conditions). Scale bars: 100, 25, and 25 pixels, respectively. b) (Boxplots) Distribution of Intersection over Union (IoU) values of StarDist models trained and tested only with WT bacteria images across multiple resolution factors (upsampling: 2^*−*2^ to 2^0^; downsampling: 2^1^ to 2^4^), showing the relationship between segmentation accuracy and the percentage of bacterial diameter covered by a single pixel. (Purple line in logarithmic scale) Theoretical throughput was calculated as the number of objects observable by a unit of time (*τ*) given a certain resolution. *τ* is specific to each microscopy acquisition modality and preserved across the different resolutions. c) Diameter distributions across resolution scales estimated from GT and the StarDist results for WT bacteria images in b). d) Comparison of the diameter distributions across various resolution scales for WT and treated bacteria. StarDist was trained with a mixed dataset containing WT and PC190723-treated *S. aureus* images in this case.

## Discussion

Our systematic investigation of the resolution paradox in deep learning-driven bioimage analysis reveals a fundamental trade-off between spatial resolution and computational performance. Our results demonstrate that popular methods like U-Net and StarDist (which often use a U-Net architecture) achieve optimal accuracy when image resolution is calibrated to match the network’s receptive field with biological object dimensions, rather than maximising resolution. This finding challenges conventional microscopy practices while offering significant practical benefits for experimental workflows.

The paradox emerges from fundamental neural network characteristics. When object features exceed the receptive field (high resolution), networks lack sufficient context for consistent boundary prediction, resulting in fragmented masks with spurious discontinuities. Conversely, excessively downsampled images (low resolution) compress critical morphological features beyond recognition. Our quantitative analysis reveals a predictable quadratic relationship between object diameter coverage per pixel and segmentation accuracy, with optimal performance for the 2D U-Net consistently occurring when pixels cover 4-16% of object diameter—irrespective of the type of specimen or imaging modality. This optimal range represents a “sweet spot” where networks balance local detail with global context.

Most significantly, we demonstrated that models trained on optimally downsampled images maintain phenotypic discrimination capabilities. In the *S. aureus* antibiotic response experiment, critical morphological differences remained detectable even at 16*×* lower resolution than commonly used (Fig. 4). This finding has profound implications for experimental design: researchers can substantially increase imaging throughput, reduce photobleaching and phototoxicity, minimise data storage requirements, and accelerate analysis pipelines—all without compromising biological insights (21, 22).

Although these principles are established in computer vision communities, they remain underappreciated in life sciences, where imaging protocols continue to emphasise the maximum achievable resolution regardless of computational analysis requirements. Our work provides a practical framework for biologists to determine optimal resolution parameters for their specific experimental contexts. By conducting preliminary resolution calibration studies before full-scale data acquisition, researchers can design more efficient imaging protocols that balance biological information content with computational performance.

The paradox persists in emerging transformer-based architectures despite their theoretical multi-scale capabilities. As demonstrated through MicroSAM (23) foundation model evaluations, fixed context windows in pre-trained models remain susceptible to resolution mismatches (Note 2, Fig. 5). However, the growing ecosystem of interactive tools like BioImage Model Zoo (6), MicroSAM (23) and DinoSim (24) creates opportunities for rapid empirical optimisation, turning this challenge into an accessible experimental design parameter.

Democratising AI in bioimaging requires deeper integration of data physics with model architectures. Current trends favour pre-trained “smart” solutions over adaptable systems, risking suboptimal application to novel experimental contexts. Future developments should prioritise dynamic acquisition systems that adjust resolution in real-time based on ongoing analysis, coupled with neural architectures explicitly encoding multi-scale relationships. By embedding these considerations into standard workflows, the field can achieve reproducible, resource-efficient microscopy pipelines that balance information content with computational sustainability.

## Methods

### Biological image datasets

Publicly available high-resolution *Caenorhabditis elegans* (*C. elegans*) minimum projection brightfield microscopy images and instance segmentation masks were obtained from (18). Publicly available *Escherichia coli* (*E. coli*) brightfield microscopy images and instance segmentation masks dataset was obtained from (19), which is available on Zenodo `10.5281/zenodo. 5550934`. In-house acquired super-resolution *Staphylococcus aureus* (*S. aureus*) cell wall structured illumination microscopy (SIM) fluorescence images and annotated instance segmentation masks dataset available on Zenodo `10.5281/ zenodo.15169017`.

### Dataset re-sampling

ReScale4DL Python library (https://github.com/HenriquesLab/ReScale4DL) was used to resize raw images and their respective masks. For *C. elegans* and *E. coli*, the instance masks were first rescaled and then, the edges and inner side were estimated using the ImageJ macro available in DeepBacs GitHub repository (19) (https://github.com/HenriquesLab/DeepBacs/). Up-sampling was computed with `catmull-rom` interpolation and nearest-neighbor interpolation methods for the raw and the mask images, respectively (all available within the NanoPyx Python package (25)). Downsampling was computed with binning and `transform.rescale` from `scikit-image` for the raw images and the masks respectively. *C. elegans* dataset was downsampled by factors 2, 2^2^, 2^3^, and 2^4^. *E. coli* was upsampled by factors 2 and 2^2^ and downsampled by factors 2 and 2^2^. *S. aureus* was upsampled by factors 2 and 2^2^ and downsampled by factors 2, 2^2^, 2^3^, and 2^4^.

### Segmentation networks

The 2D Multilabel U-Net and 2D StarDist (9) ZeroCostDL4Mic (2) notebooks were used with DL4MicEverywhere (11) with the hyperparameter configuration in Table 1.

### Evaluation

The Intersection over Union (IoU) scores visualised in Fig.s 3 and 4 were computed for the results of all the models specified in Table 1, and for the binary segmentation (*C. elegans* and *E. coli*), semantic segmentation (*C. elegans* and *E. coli*) and instance segmentation (*S. aureus*) results. For each object *j* in an image *i*, the diameter *D*_*ij*_ of non-rounded shapes (*C. elegans* and *E. coli* datasets) was computed as the median of the Euclidean Distance Transform (EDT) values contained in the object’s skeleton. For *S. aureus* dataset, *D*_*ij*_ of each bacteria was estimated by solving the equation of the circle area, given as

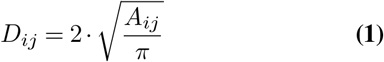

where *A*_*ij*_ is the area of the object *j* in the image *i*, measured as the sum of all the pixels in the object. To compute the average portion of diameter covered by one pixel, we first computed the average diameter for each scaling factor, *D*_*s*_ as

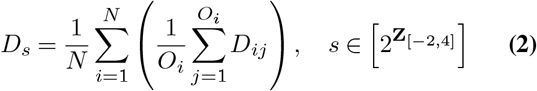

where *O*_*i*_ is the total number of objects in the image *i, N* is the total number of images in the dataset and *s* is the rescaling factor. Then, *D*_*s*_ was inverted and multiplied by 100 to compute the portion of diameter covered by one pixel.

To measure the throughput achievable for each rescaling factor *s*, we first estimated the average area in pixels of each object in an image in pixels (*A*_*s*_). Then, we consider the size of the achievable field of view (FOV) for a camera as the area in pixels of the images in the original dataset (*CAM*_*FOV*_). Considering the throughput as the amount of information recordable per unit of time, we computed the throughout as

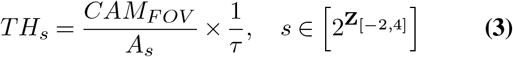

where *τ* is the time required to acquire a *FOV*. In other words, *TH*_*s*_ provides an estimate of the number of objects that can be captured by a unit of time, yet, using Equation 3 with preliminary information about the sample’s morphology and microscopy camera features, it is possible to compute this value easily.

## Code availability

The code for rescaling the images and computing the metrics is available athttps://github.com/HenriquesLab/ReScale4DL. The notebooks to train, test and deploy inference are available through the DL4MicEvrywhere (https://github.com/HenriquesLab/DL4MicEverywhere) platform.

## Data availability

WT and antibiotic-treated super-resolution *Staphylococcus aureus* (*S. aureus*) cell wall SIM images together with instance segmentation annotations are available on Zenodo https://zenodo.org/records/15169018.

## ACKNOWLEDGEMENTS

M.G.F., B.S., E.G.M., and R.H. acknowledge the support of the Gulbenkian Foundation (Fundação Calouste Gulbenkian), the European Research Council (ERC) under the European Union’s Horizon 2020 research and innovation programme (grant agreement No. 101001332) (to R.H.) and funding from the European Union through the Horizon Europe program (AI4LIFE project with grant agreement 101057970-AI4LIFE and RT-SuperES project with grant agreement 101099654-RTSuperES to R.H.). Funded by the European Union. However, the views and opinions expressed are those of the authors only and do not necessarily reflect those of the European Union. Neither the European Union nor the granting authority can be held responsible for them. This work was also supported by a European Molecular Biology Organization (EMBO) installation grant (EMBO-2020-IG-4734 to R.H.), an EMBO postdoctoral fellowship (EMBO ALTF 174-2022 to E.G.M.), a Chan Zuckerberg Initiative Visual Proteomics Grant (vpi-0000000044 with https://doi.org/10.37921/743590vtudfp to R.H.) and a Chan Zuckerberg Initiative Essential Open Source Software for Science (EOSS6-0000000260). E.G.M. acknowledges funding by Fundação para a Ciência e Tecnologia, Portugal (2023.09182.CEECIND/CP2854/CT0004). A.D.B. acknowledges the FCT 2021.06849.BD fellowship.

## Supplementary Note 1: Deep learning model training hyperparameters

**Table 1.** The network architecture and the training parameters for each dataset were preserved. The patch size for the input during training was dynamically changed with the image resizing to maintain the same field of view, except for the upsampling due to limited GPU capabilities. The values 2^*−*2,*−*1^ correspond to image upsamplings, 2^0^ is the original image resolution and 2^1,…,4^ correspond to image downsampling.

| *C. elegans* |  | *E. coli* |  | *S. aureus* |  |
| --- | --- | --- | --- | --- | --- |
| Network | 2D U-Net | Network | 2D U-Net | Network | 2D StarDist |
| Epochs | 1000 | Epochs | 1000 | Epochs | 1000 |
| Patch size<br>(per resizing<br>factor) | 512 ($\times 2^{-2, -1, 0}$)<br>256 ($\times 2$)<br>128 ($\times 2^2$)<br>64 ($\times 2^3$)<br>32 ($\times 2^4$) | Patch size<br>(per resizing<br>factor) | 512 ($\times 2^{-2, -1, 0}$)<br>256 ($\times 2$)<br>128 ($\times 2^2$) | Patch size<br>(per resizing<br>factor) | 2048 ($\times 2^{-2, -1, 0}$)<br>1184 ($\times 2$)<br>576 ($\times 2^2$)<br>272 ($\times 2^3$)<br>128 ($\times 2^4$) |
| Batch size | 5 | Batch size | 5 | Batch size | 2 |
| Validation [%] | 10 | Validation [%] | 10 | Validation [%] | 10 |
| Learning rate | 0.0001 | Learning rate | 0.0001 | Learning rate | 0.0003 |
| Pooling steps | 2 | Pooling steps | 2 | Grid parameter | 2 |
| Minimal fraction | 0.05 | Minimal fraction | 0.05 | Number of rays | 32 |

## Supplementary Note 2: Optimising the pixel size for foundation models

Here we provide a visual example of segmentation performance using foundation models for images with different pixel sizes. We performed instance segmentation of *Escherichia coli* (*E. coli*), *Caenorhabditis elegans* (*C. elegans*), and *Staphylococcus aureus* (*S. aureus*) samples using MicroSAM (23) software, with the same rescaling factors used for the Figs. 2 and 4. For simplicity, we used their 2D Annotator plugin in Napari to deploy interactive segmentation with a single-point prompt-based approach, followed by the automatic instance segmentation. Rescaled images were individually segmented by manually pointing to the same structural landmark each time. For all experiments, we chose the Vision Transformer (ViT) Large backbone architecture, a fine-tuned version of the original Segment Anything Model (SAM) (26) for cellular and nuclear segmentation in light microscopy (`ViT-l-lm`). Importantly, the segmentation results provided here may not be optimal and only aim to show the variability of the model according to the image resolution under the same conditions. Indeed, the results could be improved by changing the prompt or by increasing their number.

**Fig. 5.**
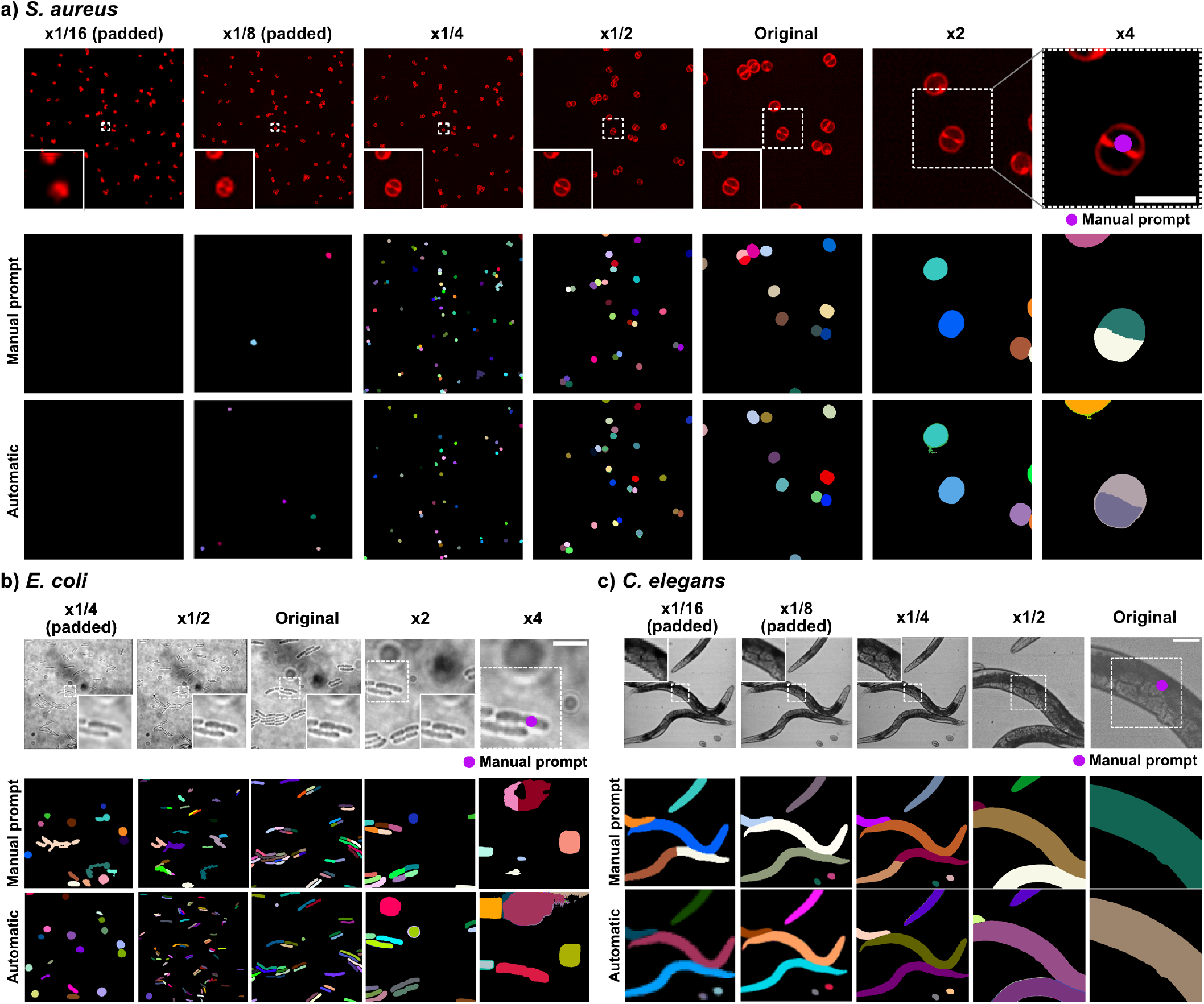
Segmentation results of Segment Anything for Microscopy on images with different pixel sizes using a single point prompting. MicroSAM (23) was used to obtain the instance segmentation of a) *S. aureus*, b) *E. coli* and c) *C. elegans* example images. For simplicity, we followed a single point-wise prompting strategy, pointing to the same object on each rescaled image (*i*.*e*., the manual prompt was the same on each panel), and run the automatic instance segmentation pipeline. All the images matched model input dimensions (512 *×* 512 pixels) to avoid internal tiling or image re-scaling. The images that had smaller dimensions were padded with zeros. Scale bars correspond to 25 *μm* for a), and 50 *μm* for b-c). Square boxes correspond to the same physical area on each image.

## Notes

https://zenodo.org/records/15169018

### Competing Interest Statement

The authors have declared no competing interest.

## Bibliography

1. Lothar Schermelleh, Alexia Ferrand, Thomas Huser, Christian Eggeling, Markus Sauer, Oliver Biehlmaier, and Gregor P. C. Drummen. Super-resolution microscopy demystified. Nature Cell Biology, 21(1):72–84, January 2019. ISSN 1476-4679. doi: 10.1038/s41556-018-0251-8. Number: 1 Publisher: Nature Publishing Group.

2. Lucas von Chamier, Romain F. Laine, Johanna Jukkala, Christoph Spahn, Daniel Krentzel, Elias Nehme, Martina Lerche, Sara Hernández-Pérez, Pieta K. Mattila, Eleni Karinou, Séamus Holden, Ahmet Can Solak, Alexander Krull, Tim-Oliver Buchholz, Martin L. Jones, Loïc A. Royer, Christophe Leterrier, Yoav Shechtman, Florian Jug, Mike Heilemann, Guillaume Jacquemet, and Ricardo Henriques. Democratising deep learning for microscopy with ZeroCostDL4Mic. Nature Communications, 12(1):2276, April 2021. ISSN 2041-1723. doi: 10.1038/s41467-021-22518-0. Publisher: Nature Publishing Group.

3. Joanna W. Pylvänäinen, Estibaliz Gómez-de Mariscal, Ricardo Henriques, and Guillaume Jacquemet. Live-cell imaging in the deep learning era. Current Opinion in Cell Biology, 85:102271, December 2023. ISSN 0955-0674. doi: 10.1016/j.ceb.2023.102271.

4. Estibaliz Gómez-de Mariscal, Carlos García-López-de Haro, Wei Ouyang, Laurène Donati, Emma Lundberg, Michael Unser, Arrate Muñoz-Barrutia, and Daniel Sage. DeepImageJ: A user-friendly environment to run deep learning models in ImageJ. Nature Methods, 18(10): 1192–1195, October 2021. ISSN 1548-7105. doi: 10.1038/s41592-021-01262-9. Publisher: Nature Publishing Group.

5. Stuart Berg, Dominik Kutra, Thorben Kroeger, Christoph N. Straehle, Bernhard X. Kausler, Carsten Haubold, Martin Schiegg, Janez Ales, Thorsten Beier, Markus Rudy, Kemal Eren, Jaime I. Cervantes, Buote Xu, Fynn Beuttenmueller, Adrian Wolny, Chong Zhang, Ullrich Koethe, Fred A. Hamprecht, and Anna Kreshuk. ilastik: interactive machine learning for (bio)image analysis. Nature Methods, 16(12):1226–1232, December 2019. ISSN 1548-7105. doi: 10.1038/s41592-019-0582-9. Publisher: Nature Publishing Group.

6. Wei Ouyang, Fynn Beuttenmueller, Estibaliz Gómez-de Mariscal, Constantin Pape, Tom Burke, Carlos Garcia-López-de Haro, Craig Russell, Lucía Moya-Sans, Cristina de-la Torre-Gutiérrez, Deborah Schmidt, Dominik Kutra, Maksim Novikov, Martin Weigert, Uwe Schmidt, Peter Bankhead, Guillaume Jacquemet, Daniel Sage, Ricardo Henriques, Arrate Muñoz-Barrutia, Emma Lundberg, Florian Jug, and Anna Kreshuk. BioImage Model Zoo: A Community-Driven Resource for Accessible Deep Learning in BioImage Analysis, June 2022. Pages: 2022.06.07.495102 Section: New Results.

7. Anastasia Mavropoulos, Chassidy Johnson, Vivian Lu, Jordan Nieto, Emilie C. Schneider, Kiran Saini, Michael L. Phelan, Linda X. Hsie, Maggie J. Wang, Janifer Cruz, Jeanette Mei, Julie J. Kim, Zhouyang Lian, Nianzhen Li, Stephane C. Boutet, Amy Y. Wong-Thai, Weibo Yu, Qing-Yi Lu, Teresa Kim, Yipeng Geng, Maddison (Mahdokht) Masaeli, Thomas D. Lee, and Jianyu Rao. Artificial Intelligence-Driven Morphology-Based Enrichment of Malignant Cells from Body Fluid. Modern Pathology, 36(8), August 2023. ISSN 0893-3952, 1530-0285. doi: 10.1016/j.modpat.2023.100195. Publisher: Elsevier.

8. Benjamin Midtvedt, Saga Helgadottir, Aykut Argun, Jesús Pineda, Daniel Midtvedt, and Giovanni Volpe. Quantitative digital microscopy with deep learning. Applied Physics Reviews, 8(1):011310, February 2021. ISSN 1931-9401. doi: 10.1063/5.0034891.

9. Uwe Schmidt, Martin Weigert, Coleman Broaddus, and Gene Myers. Cell detection with star-convex polygons. In Medical Image Computing and Computer Assisted Intervention - MICCAI 2018 - 21st International Conference, Granada, Spain, September 16-20, 2018, Proceedings, Part II, pages 265–273, 2018. doi: 10.1007/978-3-030-00934-2_30.

10. Daniel Franco-Barranco, Jesús A. Andrés-San Román, Ivan Hidalgo-Cenalmor, Lenka Backová, Aitor González-Marfil, Clément Caporal, Anatole Chessel, Pedro Gómez-Gálvez, Luis M. Escudero, Donglai Wei, Arrate Muñoz-Barrutia, and Ignacio Arganda-Carreras. BiaPy: Accessible deep learning on bioimages, January 2025. Pages: 2024.02.03.576026 Section: New Results.

11. Iván Hidalgo-Cenalmor, Joanna W. Pylvänäinen, Mariana G. Ferreira, Craig T. Russell, Alon Saguy, Ignacio Arganda-Carreras, Yoav Shechtman, Guillaume Jacquemet, Ricardo Henriques, and Estibaliz Gómez-de Mariscal. DL4MicEverywhere: deep learning for microscopy made flexible, shareable and reproducible. Nature Methods, 21(6):925–927, June 2024. ISSN 1548-7105. doi: 10.1038/s41592-024-02295-6. Publisher: Nature Publishing Group.

12. Damian Dalle Nogare, Matthew Hartley, Joran Deschamps, Jan Ellenberg, and Florian Jug. Using AI in bioimage analysis to elevate the rate of scientific discovery as a community. Nature Methods, 20(7):973–975, July 2023. ISSN 1548-7105. doi: 10.1038/s41592-023-01929-5. Publisher: Nature Publishing Group.

13. Christopher Schmied, Michael S. Nelson, Sergiy Avilov, Gert-Jan Bakker, Cristina Bertocchi, Johanna Bischof, Ulrike Boehm, Jan Brocher, Mariana T. Carvalho, Catalin Chiritescu, Jana Christopher, Beth A. Cimini, Eduardo Conde-Sousa, Michael Ebner, Rupert Ecker, Kevin Eliceiri, Julia Fernandez-Rodriguez, Nathalie Gaudreault, Laurent Gelman, David Grunwald, Tingting Gu, Nadia Halidi, Mathias Hammer, Matthew Hartley, Marie Held, Florian Jug, Varun Kapoor, Ayse Aslihan Koksoy, Judith Lacoste, Sylvia Le Dévédec, Sylvie Le Guyader, Penghuan Liu, Gabriel G. Martins, Aastha Mathur, Kota Miura, Paula Montero Llopis, Roland Nitschke, Alison North, Adam C. Parslow, Alex Payne-Dwyer, Laure Plantard, Rizwan Ali, Britta Schroth-Diez, Lucas Schütz, Ryan T. Scott, Arne Seitz, Olaf Selchow, Ved P. Sharma, Martin Spitaler, Sathya Srinivasan, Caterina Strambio-De-Castillia, Douglas Taatjes, Christian Tischer, and Helena Klara Jambor. Community-developed checklists for publishing images and image analyses. Nature Methods, 21(2):170–181, February 2024. ISSN 1548-7105. doi: 10.1038/s41592-023-01987-9. Publisher: Nature Publishing Group.

14. Virginie Uhlmann, Laurène Donati, and Daniel Sage. A Practical Guide to Supervised Deep Learning for Bioimage Analysis: Challenges and good practices. IEEE Signal Processing Magazine, 39(2):73–86, March 2022. ISSN 1558-0792. doi: 10.1109/MSP.2021.3123589. Conference Name: IEEE Signal Processing Magazine.

15. Siân Culley, Alicia Cuber Caballero, Jemima J Burden, and Virginie Uhlmann. Made to measure: An introduction to quantifying microscopy data in the life sciences. Journal of Microscopy, 295(1):61–82, 2024. ISSN 1365-2818. doi: 10.1111/jmi.13208. _eprint: https://onlinelibrary.wiley.com/doi/pdf/10.1111/jmi.13208.

16. Carsen Stringer, Tim Wang, Michalis Michaelos, and Marius Pachitariu. Cellpose: a generalist algorithm for cellular segmentation. Nature Methods, 18(1):100–106, January 2021. ISSN 1548-7105. doi: 10.1038/s41592-020-01018-x. Publisher: Nature Publishing Group.

17. Olaf Ronneberger, Philipp Fischer, and Thomas Brox. U-Net: Convolutional Networks for Biomedical Image Segmentation. In Nassir Navab, Joachim Hornegger, William M. Wells, and Alejandro F. Frangi, editors, Medical Image Computing and Computer-Assisted Intervention – MICCAI 2015, pages 234–241, Cham, 2015. Springer International Publishing. ISBN 978-3-319-24574-4. doi: 10.1007/978-3-319-24574-4_28.

18. Kevin J. Cutler, Carsen Stringer, Teresa W. Lo, Luca Rappez, Nicholas Stroustrup, S. Brook Peterson, Paul A. Wiggins, and Joseph D. Mougous. Omnipose: a high-precision morphology-independent solution for bacterial cell segmentation. Nature Methods, 19(11): 1438–1448, November 2022. ISSN 1548-7105. doi: 10.1038/s41592-022-01639-4. Publisher: Nature Publishing Group.

19. Christoph Spahn, Estibaliz Gómez-de Mariscal, Romain F. Laine, Pedro M. Pereira, Lucas von Chamier, Mia Conduit, Mariana G. Pinho, Guillaume Jacquemet, Séamus Holden, Mike Heilemann, and Ricardo Henriques. DeepBacs for multi-task bacterial image analysis using open-source deep learning approaches. Communications Biology, 5(1):1–18, July 2022. ISSN 2399-3642. doi: 10.1038/s42003-022-03634-z. Publisher: Nature Publishing Group.

20. David J. Haydon, Neil R. Stokes, Rebecca Ure, Greta Galbraith, James M. Bennett, David R. Brown, Patrick J. Baker, Vladimir V. Barynin, David W. Rice, Sveta E. Sedelnikova, Jonathan R. Heal, Joseph M. Sheridan, Sachin T. Aiwale, Pramod K. Chauhan, Anil Srivastava, Amit Taneja, Ian Collins, Jeff Errington, and Lloyd G. Czaplewski. An Inhibitor of FtsZ with Potent and Selective Anti-Staphylococcal Activity. Science, 321(5896):1673– 1675, September 2008. doi: 10.1126/science.1159961. Publisher: American Association for the Advancement of Science.

21. Guillaume Jacquemet, Alexandre F. Carisey, Hellyeh Hamidi, Ricardo Henriques, and Christophe Leterrier. The cell biologist’s guide to super-resolution microscopy. Journal of Cell Science, 133(11), June 2020. ISSN 14779137. doi: 10.1242/JCS.240713.

22. Estibaliz Gómez-de Mariscal, Mario Del Rosario, Joanna W. Pylvänäinen, Guillaume Jacquemet, and Ricardo Henriques. Harnessing artificial intelligence to reduce phototoxicity in live imaging. Journal of Cell Science, 137(3):jcs261545, February 2024. ISSN 0021-9533, 1477-9137. doi: 10.1242/jcs.261545.

23. Anwai Archit, Luca Freckmann, Sushmita Nair, Nabeel Khalid, Paul Hilt, Vikas Rajashekar, Marei Freitag, Carolin Teuber, Genevieve Buckley, Sebastian von Haaren, Sagnik Gupta, Andreas Dengel, Sheraz Ahmed, and Constantin Pape. Segment Anything for Microscopy. Nature Methods, 22(3):579–591, March 2025. ISSN 1548-7105. doi: 10.1038/s41592-024-02580-4. Publisher: Nature Publishing Group.

24. Aitor González-Marfil, Estibaliz Gómez-de Mariscal, and Ignacio Arganda-Carreras. DINOSim: Zero-Shot Object Detection and Semantic Segmentation on Electron Microscopy Images, March 2025. Pages: 2025.03.09.642092 Section: New Results.

25. Bruno M. Saraiva, Inês Cunha, António D. Brito, Gautier Follain, Raquel Portela, Robert Haase, Pedro M. Pereira, Guillaume Jacquemet, and Ricardo Henriques. Efficiently accelerated bioimage analysis with NanoPyx, a Liquid Engine-powered Python framework. Nature Methods, 22(2):283–286, February 2025. ISSN 1548-7105. doi: 10.1038/s41592-024-02562-6. Publisher: Nature Publishing Group.

26. Alexander Kirillov, Eric Mintun, Nikhila Ravi, Hanzi Mao, Chloe Rolland, Laura Gustafson, Tete Xiao, Spencer Whitehead, Alexander C. Berg, Wan-Yen Lo, Piotr Dollár, and Ross Girshick. Segment Anything. April 2023.

